# Genetic adaptation to ammonium sustains wheat grain quality and alleviates acclimation to CO_2_ enrichment

**DOI:** 10.1101/2023.11.21.568194

**Authors:** Pornpipat Kasemsap, Arnold J. Bloom

## Abstract

Plants synthesize protein through assimilating inorganic nitrogen. Yet, the extent to which soil nitrogen sources alter crop responses to atmospheric CO_2_ remains uncertain. We assessed wheat (*Triticum aestivum* L.) biomass under CO_2_ enrichment in genotypes that demonstrated a preference for ammonium (NH_4_^+^) or nitrate (NO_3_^−^), and contrasting degrees of NH_4_^+^ tolerance. Nitrogen-form preference, but not NH_4_^+^ tolerance, correlated with CO_2_ responses. Notably, NH_4_^+^-preferring genotypes maintained higher biomass and sustained grain nitrogen concentrations, thus avoiding CO_2_ acclimation, the decline in biomass stimulation after prolonged exposure to CO_2_ enrichment. Furthermore, NH_4_^+^ nutrition accelerated flowering and increased spike biomass. Breeding for NH_4_^+^-adapted genotypes may not only improve climate resilience, but also potentially accelerate development and increase yield without any penalty on grain quality. Because wheat provides 20% of the protein and carbohydrate in the human diet, our study provided strategies to sustain food security under the atmospheric conditions anticipated in the future.

**Highlight:** Breeding for NH_4_^+^-adapted genotypes may not only improve climate resilience, but also potentially accelerate development and increase yield without any penalty on grain quality under elevated CO_2_ atmospheres.

## Introduction

Anthropogenic climate change poses significant challenges to food security. Crop nutritional values decline under CO_2_ enrichment (Al-Hadeethi et al., 2019; Loladze, 2014). For example, wheat (*Triticum aestivum* L.), the staple crop that supplies 20% of protein and calories in the human diet (FAO, 2022), suffers from sharp declines in grain protein, Fe, and Zn under elevated CO_2_ concentrations (Myers et al., 2014). Maintaining crop nutritional qualities as climate changes is vital to global food security.

Declining grain protein under CO_2_ enrichment does not simply derive from dilution of organic nitrogen by additional biomass: in non-leguminous C_3_ plants, CO_2_ enrichment decreased nitrogen concentration about twice as much as the concentrations of other elements and almost three times more than the increase in carbon (Gifford et al., 2000; Loladze, 2014; Myers et al., 2014; Seibert et al., 2021; Taub et al., 2008; Uddling et al., 2018). Plants in which CO_2_ enrichment did not stimulate growth still had lower nitrogen concentrations (Broberg et al., 2017; Feng et al., 2015; Wujeska-Klause et al., 2019). Even when total nitrogen remained high under CO_2_ enrichment because of unassimilated NO_3_^−^, grain protein declined (Bloom et al., 2002). Wheat grown in California during the past 35 years compromised biomass to avoid changes in protein concentration as atmospheric CO_2_ levels increased (Bloom & Plant, 2021). Farmers who already substantially fertilize their wheat (Raun & Johnson, 1999) may need unsustainable quantities of fertilizers to maintain high yields (Pleijel et al., 2019).

Wheat responses to CO_2_ depend on nitrogen source (Bloom, 2015; Rubio-Asensio & Bloom, 2016). Crops acquire soil nitrogen primarily in the inorganic forms nitrate (NO_3_^−^) and ammonium (NH_4_^+^) (Xu et al., 2012). Elevated CO_2_ concentrations during the day (Bloom et al., 2002, 2010, 2012, 2014) and night (Asensio et al., 2015) inhibit the assimilation of NO_3_^−^ into protein in shoots, but NH_4_^+^ assimilation remains unaffected (Bloom et al., 2002). Reliance on NO_3_^−^ as the primary nitrogen source diminishes grain protein under CO_2_ enrichment in controlled environments (Bloom et al., 2010) and in the field (Bloom et al., 2014). By contrast, reliance on NH_4_^+^ sustains grain protein at elevated CO_2_ levels (Fernando et al., 2017). Breeding cultivars for a greater reliance on NH_4_^+^ might ensure future productivity (Bloom et al., 2010, 2014).

Exposure to high NH_4_^+^ concentrations, however, can be toxic (Britto & Kronzucker, 2002). Plants expend up to 40% of root respiration to prevent excessive NH_4_^+^ accumulation (Britto et al., 2001). NH_4_^+^-sensitive plants experience physiological and morphological disorders that limit growth and development (Esteban et al., 2016). To avoid toxicity, plants assimilate most of the absorbed soil NH_4_^+^ into organic compounds in roots and transport them to other organs (Britto & Kronzucker, 2002). Sensitivity to nitrogen forms, especially exposure to high NH_4_^+^ concentrations, varies significantly among species (Bloom et al., 2012; Britto & Kronzucker, 2013; Miller & Cramer, 2005). Nevertheless, information on the genetic basis of such responses remains sparse.

Rarely do studies account for different inorganic nitrogen forms (Kasemsap & Bloom, 2022). Only a handful of studies have contrasted the physiological responses of wheat to NH_4_^+^ versus NO_3_^−^ and CO_2_ levels (Bloom et al., 2002; Carlisle et al., 2012; Rubio-Asensio & Bloom, 2016), and none of the studies selected the plant materials based primarily on the degree of preference for a nitrogen form or NH_4_^+^ tolerance. The limited genetic diversity in any given study (Kasemsap & Bloom, 2022) and the difficulties in precisely controlling the nitrogen forms in the root zone (Bloom et al., 2012) may underlie the lack of understanding about this topic.

Here, we characterized biomass under CO_2_ enrichment of 12 genotypes from diverse mapping populations that demonstrated a preference for NH_4_^+^ or NO_3_^−^ and high tolerance or susceptibility to NH_4_^+^ toxicity. Given the strong interactions between carbon and nitrogen metabolisms, we hypothesized that plants would be more responsive to their preferred nitrogen form under CO_2_ enrichment. These traits can guide breeders in selecting genotypes that ensure grain yield and quality under the edaphic and atmospheric conditions anticipated in the near future.

## Materials and methods

### Plant materials

In a preliminary study, we assessed the biomass of 15-day (d) old hexaploid wheat plants under either NH_4_^+^ or NO_3_^−^ nutrition at ambient CO_2_ concentrations and calculated the ratio of plant biomass under 1) 20 mM vs. 0.5 mM NH_4_^+^ (NH_4_^+^ tolerance) and 2) 0.5 mM NO_3_^−^ vs. 0.5 mM NH_4_^+^ (nitrogen-form preference). Here, based on the nitrogen responses, we selected 3 genotypes from a panel of 875 spring hexaploid wheat (*Triticum aestivum* L.) accessions from the USDA-ARS National Small Grain Core Collection (NSGCC) (Maccaferri et al., 2015) and 8 genotypes from 2 bi-parental Nested-Association Mapping (NAM) populations (Blake et al., 2019; Zhang et al., 2018). We included two additional genotypes: cv. Veery 10, which we have used in previous studies, as a reference genotype (Asensio et al., 2015; Bloom et al., 2002, 2012, 2020; Carlisle et al., 2012; Rubio-Asensio & Bloom, 2016), and cv. Berkut, the common parent of the bi-parental populations.

We defined inorganic nitrogen-form preference for each genotype as “NH_4_^+^-preference”, “NO_3_^−^ -preference”, or potentially adapted to “both forms” when the biomass ratio under 0.5 mM NH_4_^+^ vs. NO_3_^−^ was higher than 1.1, lower than 0.9, or between 0.9 – 1.1, respectively. We defined NH_4_^+^ tolerance for each genotype as either “tolerant” or “susceptible” when the biomass ratio of plants under 20 mM vs. 0.5 mM NH_4_^+^ was higher or lower than 0.9, respectively. Based on these criteria, our experimental panel consisted of one reference genotype, one common parent of the bi-parental mapping populations and 11 additional genotypes for which we had sufficient seeds and that showed diverse responses to the nitrogen source (Table 1).

**Table 1.** Selected genotypes based on the ratio of 15 d plant biomass at ambient CO_2_ levels under 1) 0.5 mM NO_3_^−^ vs. 0.5 mM NH_4_^+^ (nitrogen-form preference) and 2) 20 mM vs. 0.5 mM NH_4_^+^ (NH_4_^+^ tolerance) from the USDA-National Small Grain Core Collection (NSGCC; 875 genotypes) and bi-parental Nested Association Mapping populations for which cv. Berkut was the common female parent (NAM; 75 genotypes each). Accession or line ID are from the original studies where the populations were developed.

| # | Population (Accession/line ID) and name used in this study |  | Biomass ratio<br>NO<sub>3</sub><sup>-</sup>: NH<sub>4</sub><sup>+</sup> | Form preference | Biomass ratio<br>High: Low NH<sub>4</sub><sup>+</sup> | NH<sub>4</sub><sup>+</sup> tolerance |
| --- | --- | --- | --- | --- | --- | --- |
| 1 | NSGCC (PI83733) | 74 | 0.96 | Both | 0.98 | Tolerant |
| 2 | NSGCC (PI182077) | 166 | 1.07 | Both | 1.02 | Tolerant |
| 3 | NSGCC (PI285944) | 459 | 1.05 | Both | 0.68 | Susceptible |
| 4 | NAM Parent | Berkut | 1.16 | NO<sub>3</sub><sup>-</sup> | 0.71 | Susceptible |
| 5 | BerkutxPBW343 (PBW343_15) | P10 | 0.71 | NH<sub>4</sub><sup>+</sup> | 0.45 | Susceptible |
| 6 | BerkutxPBW343 (PBW343_40) | P22 | 1.19 | NO<sub>3</sub><sup>-</sup> | 1.12 | Tolerant |
| 7 | BerkutxPBW343 (PBW343_52) | P33 | 0.78 | NH<sub>4</sub><sup>+</sup> | 1.13 | Tolerant |
| 8 | BerkutxPBW343 (PBW343_58) | P39 | 1.24 | NO<sub>3</sub><sup>-</sup> | 0.48 | Susceptible |
| 9 | BerkutxRAC875 (RAC875_45) | R33 | 0.58 | NH<sub>4</sub><sup>+</sup> | 1.02 | Tolerant |
| 10 | BerkutxRAC875 (RAC875_57) | R39 | 0.82 | NH<sub>4</sub><sup>+</sup> | 0.58 | Susceptible |
| 11 | BerkutxRAC875 (RAC875_70) | R49 | 1.33 | NO<sub>3</sub><sup>-</sup> | 0.94 | Tolerant |
| 12 | BerkutxRAC875 (RAC875_99) | R69 | 0.78 | NH<sub>4</sub><sup>+</sup> | 0.50 | Susceptible |
| 13 | Reference | Veery 10 | 1.03 | Both | 0.75 | Susceptible |

### Growth conditions

For each genotype, we used seeds from the same mother plant grown in a controlled environmental chamber at Davis, California, USA. We disinfected seeds with 10% v/v NaHClO_3_ for 10 minutes, rinsed them multiple times with deionized water, and placed them on a germination paper soaked with deionized water. We kept the seeds on the germination paper at 4°C for at least 48 hr and later moved them to room temperature for 4 d until transplanting. We transplanted uniform seedlings into 20 dm^3^ opaque tubs containing an aerated nutrient solution composed of 1 mM K_2_HPO_4_, 1 mM KH_2_PO_4_, 2 mM MgSO_4_, 2 mM CaCl_2_, 1 ppm Fe-NaDTPA (Sequestrene 330, Becker Underwood), and 50% of other micronutrients (B, Mn, Zn, Cu, Mo) in a Hoagland solution (Epstein & Bloom, 2005) with custom nitrogen concentrations. We adjusted the solution pH in the tubs initially to 6.0 and replenished the tubs with freshly made solution twice in the first week and every 1 to 2 days in the following weeks to maintain pH and sufficient nutrient levels. The tubs were kept in controlled environmental chambers under 16 h of 900 µmol m^-2^ s^-1^ of light at plant height and 22/15°C and 60/83 % relative humidity during the light and dark period respectively (Conviron, Winnipeg, Canada). We passed industrial CO_2_ gas from a compressed gas tank through a column filled with KMnO_4_-covered clay pebbles to remove any C_2_H_2_ contaminants and injected it into the growth chambers to regulate the CO_2_ concentration of two controlled environment chambers at either ∼420 ppm (“ambient”) or 750 ppm as monitored by non-dispersive infrared analyzers.

We used a completely randomized design (Supplementary Figure S1) to evaluate the influence of CO_2_ levels and inorganic nitrogen forms on the growth of the 13 genotypes. We placed a group of 12 tubs into each of the two growth chambers. Six tubs in each chamber received either 0.5 mM NH_4_^+^ from (NH_4_)_2_SO_4_ or 0.5 mM NO_3_^−^ from KNO_3_ as a sole nitrogen source. Each tub represented an experimental unit and contained 42 plants: 3 seedlings of each of the 12 genotypes and 6 seedlings of the control genotype (cv Veery 10) placed in the middle of the tub. Each group of seedlings of the same genotype was randomly transplanted into the tub. We rotated the tubs regularly within each chamber to minimize positional effects. The randomization of genotypes within each tub should have alleviated shading among heterogeneous canopy architectures.

### Biomass assessment

To determine biomass production, we harvested one plant of each genotype from each tub at 21 d during the tillering stage and at 60 d after transplanting when most genotypes had completed the heading stage. We took 6 biological replicates per genotype per nitrogen treatment, one plant from each tub, in each destructive harvest. We harvested 2 and 4 plants of Veery 10 at each harvest, respectively. As we harvested plants to measure growth, the planting density changed from 217 plants m^-2^ (42 plants per a 20” × 15” tub surface) at the beginning of the experiment to 145 plants m^-2^ (28 plants per tub) after the first harvest. Planting density in this experiment remained within the range of other controlled environmental studies as well as the field settings of 80-400 plant m^-2^ (Fischer et al., 2019). We harvested the roots and shoots separately during the first harvest. Because the roots became entangled after 3 weeks of growth, we assessed only shoot biomass at the second harvest. We rinsed plant samples with deionize water and dried them at 60°C for at least 48 h before weighing them.

To quantify changes in yield components, we determined tiller number, tiller producing spike number, spike number, spike biomass, spikelet number and grain nitrogen content as proxies for potential grain yield. Spike biomass refers to dry mass of all harvested spikes, including both chaffs and grains. We did not measure mass of individual spike components, because the kernels were not fully developed across all genotype and nitrogen treatments. We selected the longest (most developed) and the shortest (least developed) spikes of each plant to determine the number of rachis and spikelets. We define spikelets as rachis whose florets developed into at least one seed. We calculated spike fertility as a ratio of spikelet number to total rachis number. Except for genotype “74”, “166”, “459”, we sampled developing grains randomly from the longest spike of each plant and measured nitrogen concentration with a Leco TrusSpec N (Leco Corporation, St. Joseph, USA) that uses a combustion method (Association of Official Analytical Chemists, 2000) coupled with an infrared detection system. The kernels of genotype “P10”, “P33” and “P39” developed slower under NO_3_^−^ than NH_4_^+^ nutrition but were large enough for the analysis. We were unable to collect data on the mature grains because our growth chamber failed and subjected the plants to temperatures above 70°C. Nonetheless, the relative differences in nitrogen concentrations of developing grains allowed us to compare nitrogen and carbon translocations to the grains among the treatments.

### Statistical modeling

We quantified the influence of nitrogen form, CO_2_ concentration, and genotype with a linear model: biomass ∼ nitrogen form × CO_2_ concentration × genotype. Because of differences in sample sizes, we analyzed biomass of the control genotype, Veery 10, separately. We conducted an analysis of variance (ANOVA) and calculated type II sums of squares (Langsrud, 2003) using R/car version 3.1 (Fox & Weisberg, 2019). We checked the ANOVA assumptions for homogeneity of variance and normal distribution of model residuals with Levene’s test (Levene, 1960) and Shapiro-Wilk Normality test (Royston, 1982), respectively. Based on ANOVA outputs, we determined mean separations with Tukey’s Honest Significant Difference (Tukey’s HSD) method in R/emmeans version 1.8.6 (Lenth, 2023) and R/multcomp version 1.4-23 (Hothorn et al., 2008).

To determine whether the classification of inorganic nitrogen-form preference and NH_4_^+^ tolerance of individual genotypes were associated with biomass production, we updated the linear regression model with the categorical variables based on the classification of nitrogen-form adaptation and tolerance, as followed: (*a*) biomass ∼ nitrogen form × CO_2_ concentration × inorganic nitrogen-form preference or (*b*) biomass ∼ nitrogen form × CO_2_ concentration × NH_4_^+^ tolerance.

### Validation of the reference genotype, Veery 10

Before we examined genotypic variations of biomass and partitioning, we verified growth and development of the reference genotype, Veery 10, a semi-dwarf cultivar commonly used for experiments (Bishop & Bugbee, 1998; Waines & Ehdaie, 2007). Studies in controlled environments have often used cv Veery 10 due to its compact structure and fast growth rate (Supplementary Table S1). We analyzed growth of Veery 10 separately from the other genotypes because this reference genotype experienced a slightly different growth environment due to the experimental design and had 2 to 4 times as many replicates. Since an early study from our group that used Veery 10 (Smart et al., 1998), atmospheric CO_2_ levels have risen from about 366 ppm to over 420 ppm (IPCC, 2021). As such, controlled experiments have also increased the “ambient” concentrations about ∼15% to keep up with natural atmospheric changes (Supplementary Table S1). Despite the increased ambient CO_2_ levels and slight differences in growth conditions across studies, we still observed consistent trends in responses to CO_2_ enrichment (Supplementary Figure S2, Supplementary Figure S3, and Supplementary Table S1). Veery 10 produced similar biomass under 0.5 mM NO_3_^−^ and 0.5 mM NH_4_^+^ at ambient CO_2_ (Supplementary Figure S2 and Supplementary Figure S3). In our previous experiments, we also observed that Veery 10 at ambient CO_2_ produced comparable biomass under different nitrogen forms (Table 1).

The influence of nitrogen form and CO_2_ enrichment, however, differed as plants transitioned into the reproductive stages. In Veery 10, during the vegetative stage at 21 d, CO_2_ enrichment increased shoot biomass 61% and 37% under NH_4_^+^ and NO_3_^−^, respectively (Supplementary Figure S2B). Veery 10 partitioned significantly more biomass to shoot growth and had a higher shoot-to-root ratio under NH_4_^+^ than NO_3_^−^ under CO_2_ enrichment, although differences in absolute shoot and whole plant mass were not statistically significant between the two nitrogen forms (Supplementary Figure S2A-E). Our results still supported that elevated CO_2_ concentrations enhanced biomass of shoots under NH_4_^+^ nutrition and roots under NO_3_^−^ nutrition, respectively (Rubio-Asensio & Bloom, 2016). Veery 10 was most responsive to changes in CO_2_ level when receiving NH_4_^+^ as a sole nitrogen source (Carlisle et al., 2012). Yet we observed a slight increase in biomass under NH_4_^+^ (Bloom et al., 2012; Rubio-Asensio & Bloom, 2016), rather than the decrease previously reported (Carlisle et al., 2012).

At the reproductive stage, we can only compare shoot growth because we could not separate roots from individual plants (Supplementary Figure S3). For Veery 10 under elevated CO_2_ exposure, the average shoot growth stimulation by CO_2_ across nitrogen forms decreased from +50% at 21 d to +18% at 60 d (Supplementary Figure S3A). Changes in CO_2_ and nitrogen form did not affect tiller and spike number but had profound consequences on relative development of Veery 10 (Supplementary Figure S3C-G). On one hand, under CO_2_ enrichment ∼25% tillers bore no spikes under NH_4_^+^ (Supplementary Figure S3G), although total spike mass remained comparable to plants under NO_3_^−^ (Supplementary Figure S3C). On the other hand, CO_2_ enrichment enhanced shoot biomass partitioning to spikes more under NO_3_^−^ than NH_4_^+^, primarily because the fraction of spike biomass at ambient CO_2_ levels under NO_3_^−^was significantly lower than under NH_4_^+^ (Supplementary Figure S3D). In the main result, we focus on the phenotypic diversity in the panel of 12 genotypes that had responses distinct from those we observed in the reference genotype Veery 10.

## Results

The genetic influence was highly significant across all traits, but its interactions varied (Supplementary Table S1). Responses of the reference genotype Veery 10 were consistent with previous studies (see Methods) but failed to represent the significant, much larger biomass variation exhibited among other genotypes (Figure 1). Both nitrogen forms had a similar influence on most genotypes at ambient CO_2_ levels, but biomass under CO_2_ enrichment varied among genotypes and between NO_3_^−^ and NH_4_^+^ nutrition (Figure 1A).

**Figure 1.**
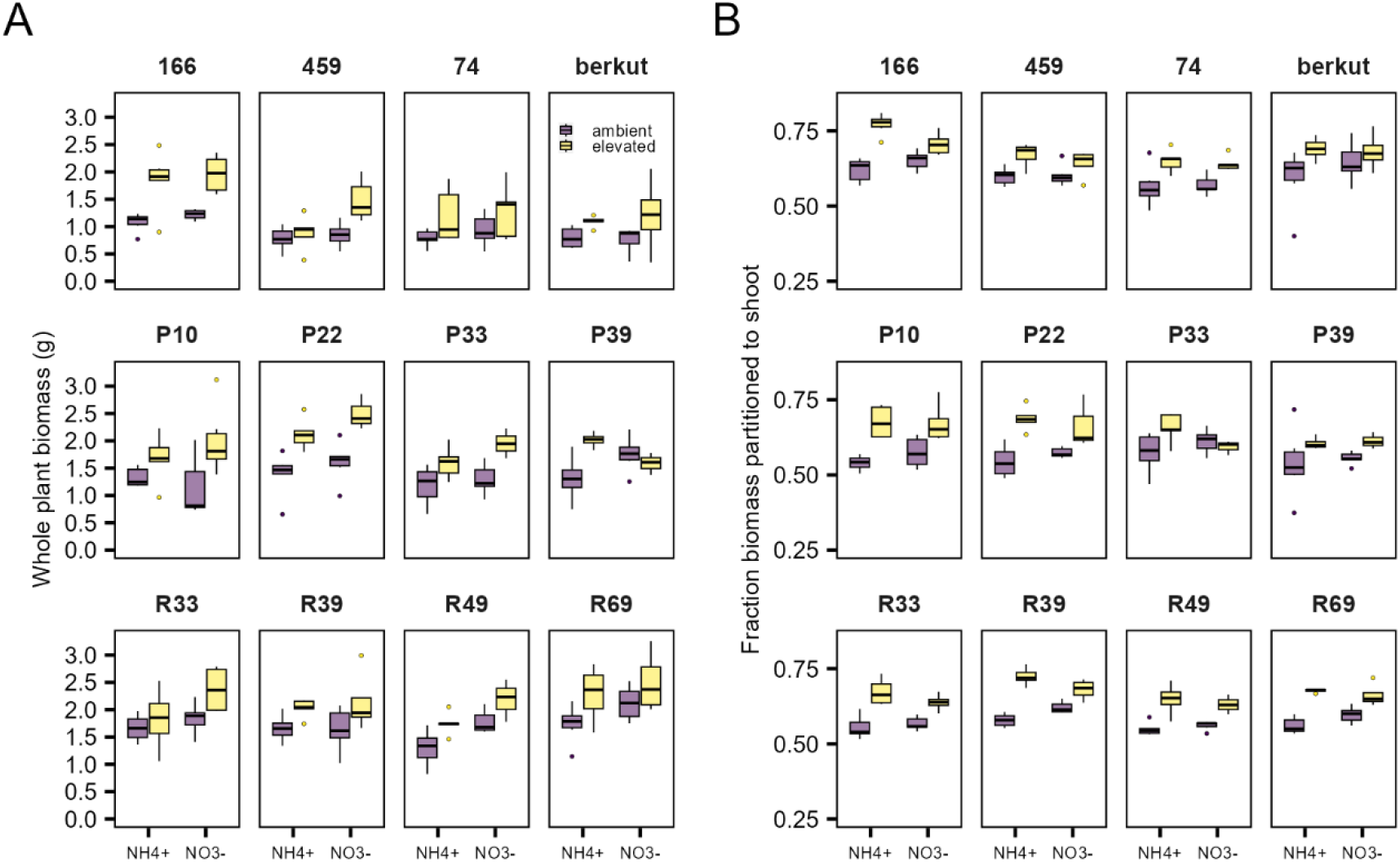
CO_2_ enrichment enhanced vegetative biomass production and favored partitioning to shoot under NH_4_^+^ nutrition. Distribution of whole plant biomass (A) or fraction biomass partitioned to root (B) of 12 wheat genotypes under NH_4_^+^ or NO_3_^−^ as a sole nitrogen source at 21 d after transplanting in ambient (purple box) or elevated (yellow box) CO_2_ concentrations. The box plots represent minima, 25th percentiles, medians, 75th percentiles and maxima (n=6). Each dot represents outliers.

### Biomass partitioning under CO_2_ enrichment depends on nitrogen source

CO_2_ enrichment favored biomass partitioning to shoots under NH_4_^+^ nutrition for 4 out of 12 genotypes, whereas the remaining genotypes partitioned similar proportions to roots and shoots under both forms (Figure 1B and Supplementary Figure S4). Genetics interacted with either nitrogen and CO_2_ in determining growth of the organ involved in their assimilation: for root biomass in response to nitrogen forms (Figure 2A) and for shoot biomass in response to CO_2_ levels (Figure 3A & B). Linear relationships between shoot and root biomass across all genotypes demonstrated that plants under CO_2_ enrichment grew more root with NO_3_^−^ nutrition and more shoot with NH_4_^+^ nutrition, and the differences increased as the plant size increased (Figure 2B).

**Figure 2.**
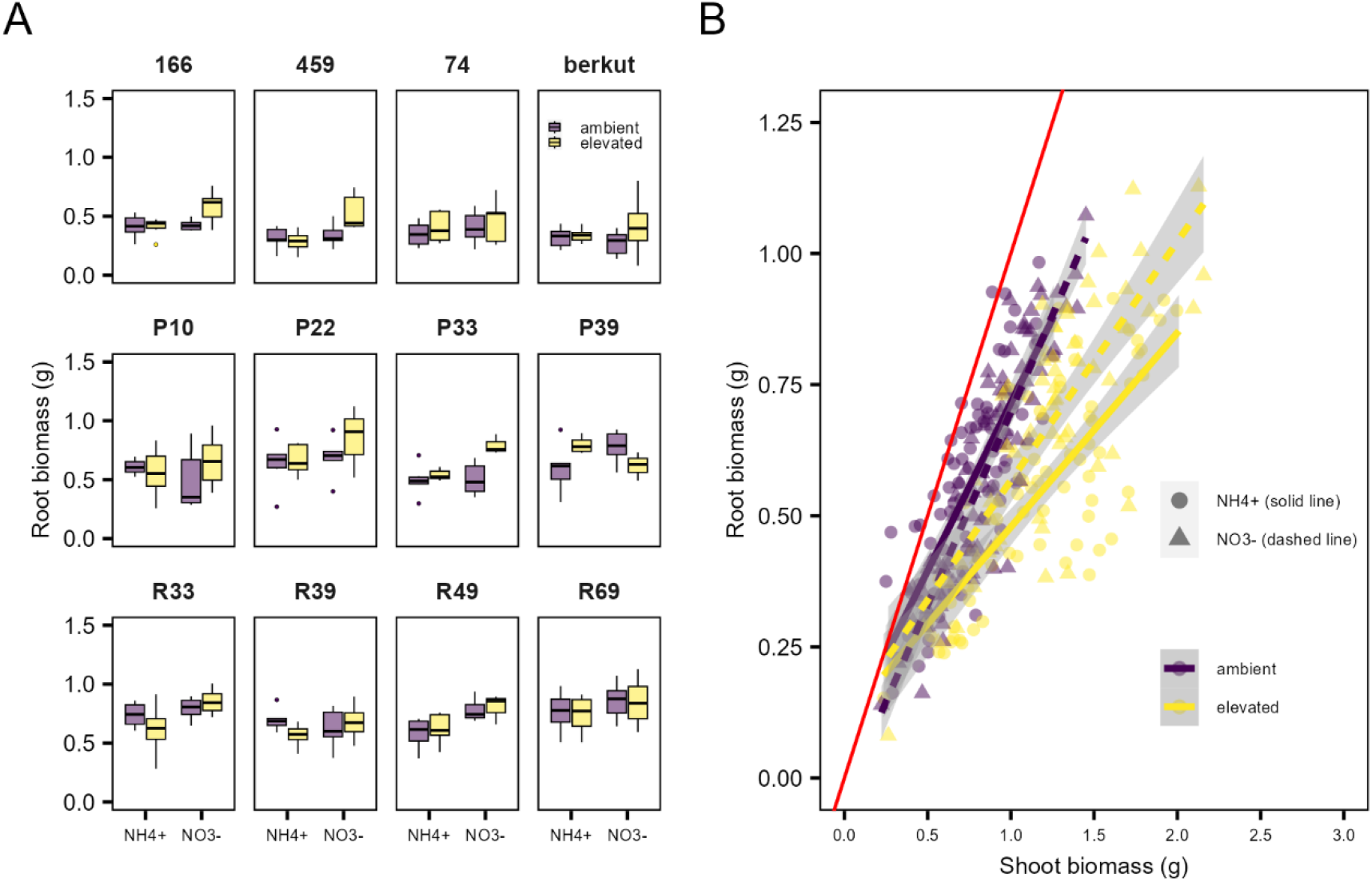
Elevated CO_2_ enhancement of root growth under NO_3_^−^ and shoot growth under NH_4_^+^ nutrition amplified with plant size. Distribution of root biomass (A) under NH_4_^+^ or NO_3_^−^ as a sole nitrogen source at 21 d after transplanting in ambient (purple box) or elevated (yellow box) CO_2_ concentrations. The box plots represent minima, 25^th^ percentiles, medians, 75^th^ percentiles and maxima (n=6). Each dot represents outliers. Scatter plots and linear relationships between shoot and root biomass across 12 wheat genotypes (B) under NH_4_^+^ (purple line) or NO_3_^−^ (yellow line) at ambient (circle, solid line) or 750 ppm CO_2_ (triangle, dashed line). The red line represents a 1:1 ratio. Gray areas represent 95% confidence intervals.

**Figure 3.**
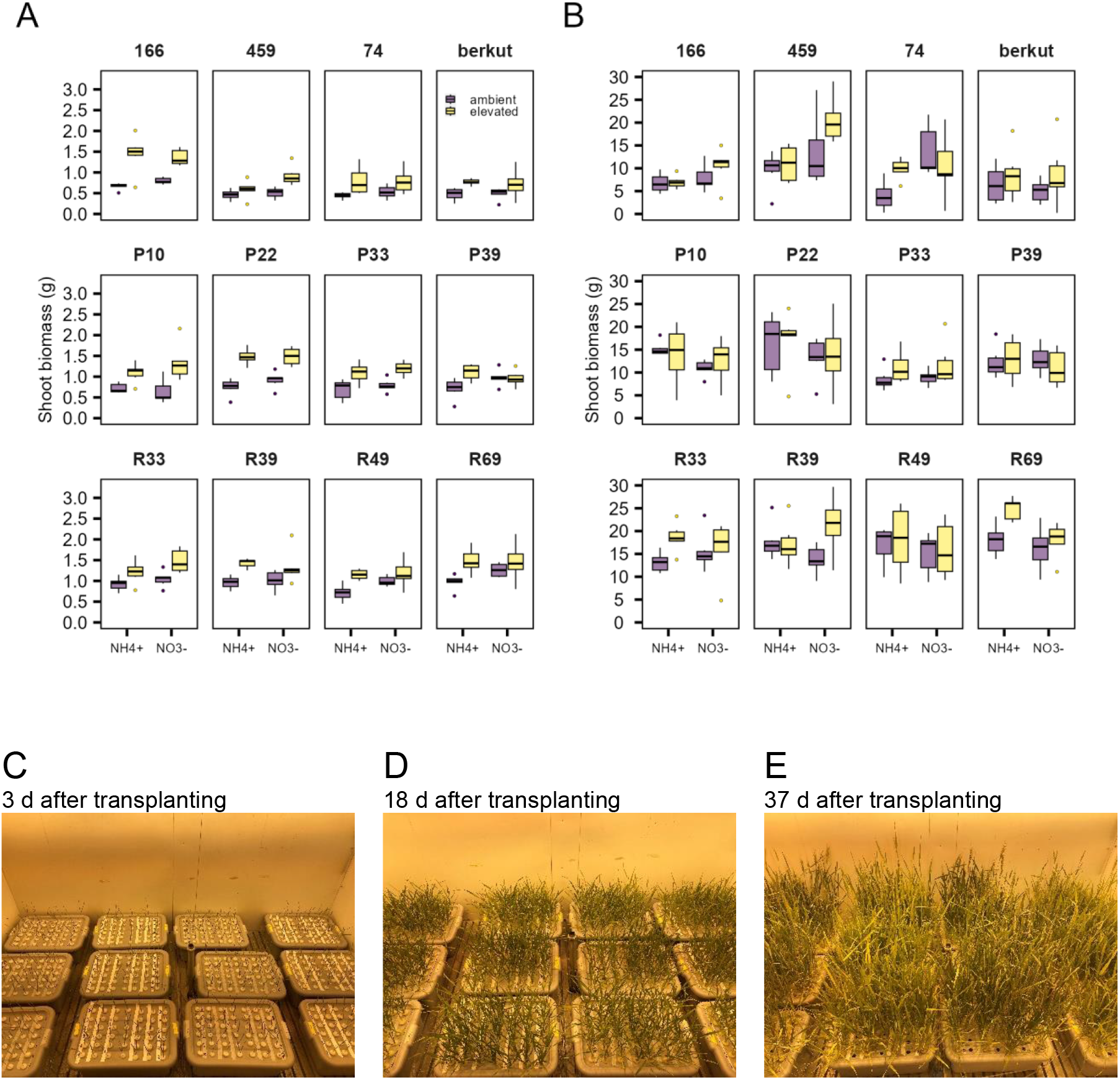
As plants transitioned into the reproductive stage, effects of CO_2_ interactions with nitrogen forms varied among genotypes. Distribution of shoot biomass at 21 d (A) and 60 d (B) after transplanting of 12 wheat genotypes under NH_4_^+^ or NO_3_^−^ as a sole nitrogen source at 21 d after transplanting in ambient (purple box) or elevated (yellow box) CO_2_ concentrations. The box plots represent minima, 25th percentiles, medians, 75th percentiles and maxima (n=6). Each dot represents outliers. Time course canopy change in the ambient CO_2_ chamber at 3, 18 and 37 d after transplanting respectively (C-E). Leaves covered most of the tub surface by 3 weeks (D), and the canopy fully closed by 5 weeks after transplanting (E).

### Genotypic variations determined CO_2_ acclimation

We focused on the influence of CO_2_ enrichment on shoot growth as the plants transitioned into reproduction (Figure 3). Genotypic variations in acclimation to CO_2_ was apparent after exposure to elevated CO_2_ for 60 d (Figure 3A-B and Supplementary Table S1). The biomass stimulation of CO_2_ diminished or even became negative (Figure 3B). For example, genotype “166”, the biomass increase under CO_2_ enrichment with NH_4_^+^ nutrition went from 117% at 21 d to negligible at 60 d, whereas with NO_3_^−^ nutrition the effects halved (71%→33%) (Figure 3A-B). On average, NH_4_^+^-grown plants were more responsive to changes in CO_2_ concentrations (Figure 3A-B): the biomass stimulation from 21 d to 60 d changed from 61% to 19% under NH_4_^+^ and 42% to 18% under NO_3_^−^. Notably, not all genotypes suffered from CO_2_ acclimation (Figure 3B); a few genotypes maintained biomass stimulation. For example, biomass stimulation increased from 31% to 41% with NH_4_^+^ nutrition for “R33” and from 32% to 55% with NO_3_^−^ nutrition for “R39” (Figure 3B).

### NH_4_^+^ accelerated reproductive development

The reference genotype, Veery 10 reached anthesis about 7 d earlier under NH_4_^+^ than under NO_3_^−^. Similarly, NH_4_^+^ nutrition accelerated the flowering of most other genotypes (Figure 4). We cannot distinguish whether final grain yield differed because we did not have complete data on the precise differences in flowering time or biomass at physiological maturity, but the growth data at 60 d provided a “snapshot” into crop development that can serve as a proxy for potential yield.

**Figure 4.**
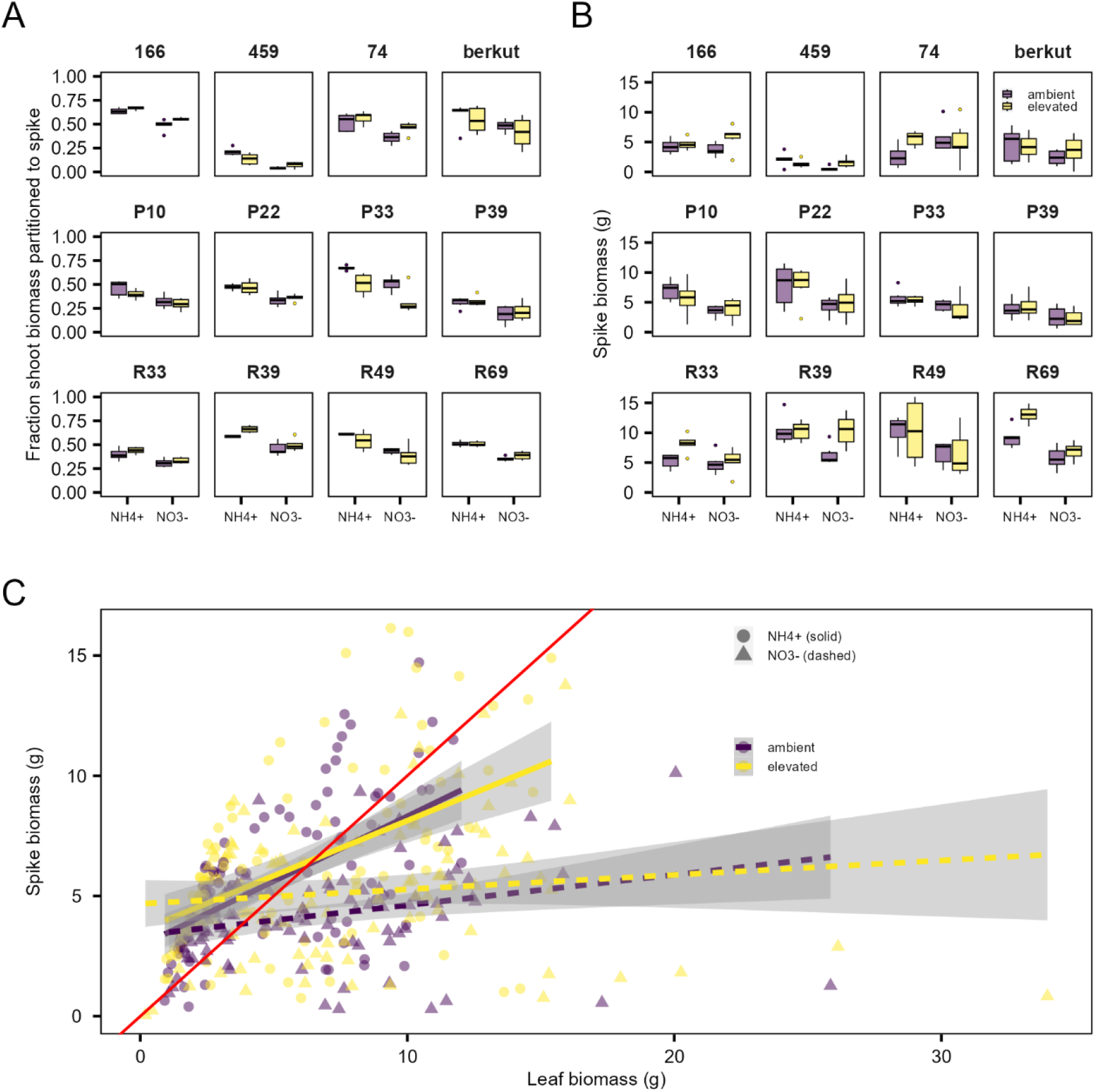
NH_4_^+^ nutrition influenced flowering and accelerated spike development. Distribution of fraction shoot biomass partitioned to spike (A), spike biomass at 60 d (B) after transplanting of 12 wheat genotypes under NH_4_^+^ or NO_3_^−^ as a sole nitrogen source at 21 d after transplanting in ambient (purple box) or elevated (yellow box) CO_2_ concentrations. The box plots represent minima, 25^th^ percentiles, medians, 75^th^ percentiles and maxima (n=6). Each dot represents outliers. Scatter plots and linear relationships between leaf and spike biomass at 60 d across 12 wheat genotypes (C) under NH_4_^+^ (purple line) or NO_3_^−^ (yellow line) at ambient (circle, solid line) versus 750 ppm CO_2_ (triangle, dashed line). The red line represents a 1:1 ratio. Gray areas represent 95% confidence intervals.

We examined how resource allocation shifted between the vegetative and the reproductive stages. Vegetative tiller development was controlled by nitrogen form and the interactions between CO_2_ level and genotype (Supplementary Table S1). Tiller numbers were slightly higher under NO_3_^−^ than under NH_4_^+^ (Supplementary Figure S5A). Because tiller buds almost fully developed by the first harvest, we observed just a few more tillers at 60 d (Supplementary Figure S5B). Similar to biomass, tiller growth enhancement by CO_2_ was slower at 60 d than at 21 d (Supplementary Figure S5A-B). Interactions of the nitrogen and genotype became more important than CO_2_ in controlling the reproductive tiller number (Supplementary Table S1). With NO_3_^−^ nutrition, 4 out of 13 genotypes had slightly lower tiller number under CO_2_ enrichment than under the ambient treatment (Supplementary Figure S5B). The number of spikes did not differ between nitrogen forms nor CO_2_ levels (Supplementary Figure S5D), although the proportion of tillers that produced spikes varied across treatments (Supplementary Figure S5C).

The influence of nitrogen was most pronounced in the allocation of shoot biomass to spikes (Figure 4A). At 60 d, plants allocated more biomass to spikes with NH_4_^+^ than with NO_3_^−^ nutrition, suggesting faster development under NH_4_^+^ nutrition (Figure 4B). CO_2_ enrichment increased absolute spike mass ∼23% across both nitrogen forms. (Figure 4B), but the influence on spike mass allocation varied with genotype. Most genotypes increased or showed no change in spike partitioning, except for “Berkut”, “P33” and “R49” (Figure 4A). The influence of nitrogen form on spike mass also varied with genotype. Most genotypes produced higher spike mass with NH_4_^+^ nutrition, with a few exceptions (Figure 4B); for example, “74” had 53% more spike mass with NO_3_^−^ than with NH_4_^+^ across the CO_2_ levels, or more than double the size if we consider only the ambient CO_2_ treatment.

We determined rachis and spikelet number in the most and least developed spike of each plant (Supplementary Figure S6). Nitrogen form influenced the rachis number and the proportion of rachis that developed into spikelets, an indicator of spike fertility (Supplementary Figure S6). Although the rachis numbers were generally higher in NO_3_^−^-grown plants (Supplementary Figure S6A-B), spikelet numbers were relatively similar under both nitrogen forms when spikes reached maturity (Supplementary Figure S6C). We observed that basal spikelets were more likely to abort. As a result, spike fertility of the most developed spike was relatively higher under NH_4_^+^ nutrition for most genotypes (Supplementary Figure S7).

Spike biomass was positively correlated with leaf biomass (Figure 4C) although there was no clear pattern on how nitrogen form and CO_2_ level influenced leaf biomass (Supplementary Figure S8). CO_2_ enrichment generally increased leaf biomass, but the extent of increase depended on nitrogen form and genotype (Supplementary Table S1). Interestingly, as leaf biomass increased, plants receiving NH_4_^+^ produced significantly higher spike mass than plants receiving NO_3_^−^ (Figure 4C). Such relationships affirm the positive influence of NH_4_^+^ on spike growth and development.

We measured nitrogen concentration of kernels in the most developed spikes (Figure 5). To limit developmental differences, we focused on the genotypes from the two bi-parental populations and their common parent Berkut because their developmental rates were somewhat similar, although spikes of genotype “P10”, “P33”, and “P39” in the NO_3_^−^ treatment still developed much slower than those in the NH_4_^+^ treatment (Figure 4B). Grain nitrogen concentration was lower in NO_3_^−^ (2.32%) than NH_4_^+^ (3.07%) and their responses to CO_2_ enrichment depended on genotype (Figure 5). Under CO_2_ enrichment, nitrogen concentrations either decreased or did not change (Figure 5). Grains of genotype “P10” and “P33” had higher nitrogen concentration under NO_3_^−^ at the elevated CO_2_ level, but such an anomalous trend likely derived from nitrogen form-driven differences in developmental stages because these kernels were much smaller than other genotypes and the same genotype under NH_4_^+^ (Figure 5). When we exclude extremely small kernels with higher concentrations from these genotypes, the data pattern of the subset genotypes remain consistent with the whole dataset (Supplementary Figure S9).

**Figure 5.**
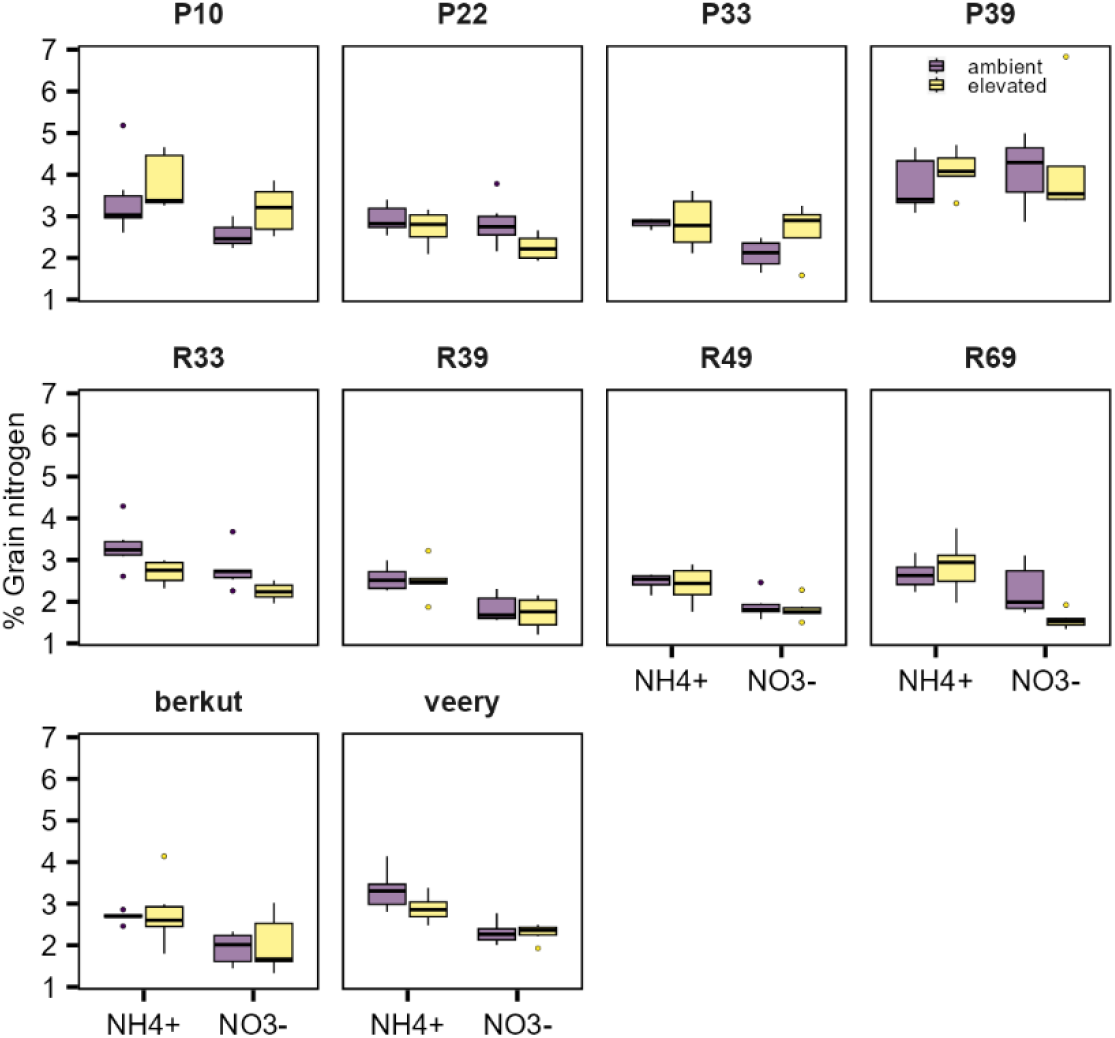
Nitrogen concentrations in developed grains were lower under NO_3_^−^ than NH_4_^+^ and dependent upon genetics and CO_2_ concentrations interactions. Distribution of nitrogen concentrations in developing grains in the most developed spike of 10 wheat genotypes under NH_4_^+^ or NO_3_^−^ as a sole nitrogen source at 60 d after transplanting in ambient (purple box) or elevated (yellow box) CO_2_ concentrations. The box plots represent minima, 25^th^ percentiles, medians, 75^th^ percentiles and maxima (n=6). Each dot represents outliers. Genotype “74”, “166” and “459” were excluded from nitrogen analysis due to significant differences in developmental stages.

### Nitrogen adaptation explained responses to CO_2_ enrichment

We examined whether nitrogen-form preference or NH_4_^+^ tolerance can explain biomass change under CO_2_ enrichment (Supplementary Table S2). Because the three genotypes from the NSGCC panel responded similarly to both nitrogen forms (Table 1), we retained only genotypes from the two bi-parental populations and their common parent that had a preference for either nitrogen form in the analysis. Nitrogen preference accounted for ∼4% of the variation in biomass (Supplementary Table S2). Genotypes that preferred NH_4_^+^ produced more whole plant biomass at 21 d (Figure 6A), shoot biomass at 60 d (Figure 6B), and spike biomass (Figure 6C), independent of nitrogen form and CO_2_ level. But, nitrogen-form preference did not alter grain nitrogen (Figure 6D). Moreover, tolerance to high NH_4_^+^ levels was not associated with biomass under the non-toxic concentrations in the current experiment except for grain nitrogen that differed slightly between tolerant and susceptible genotypes (P=0.046) (Supplementary Figure S10).

**Figure 6.**
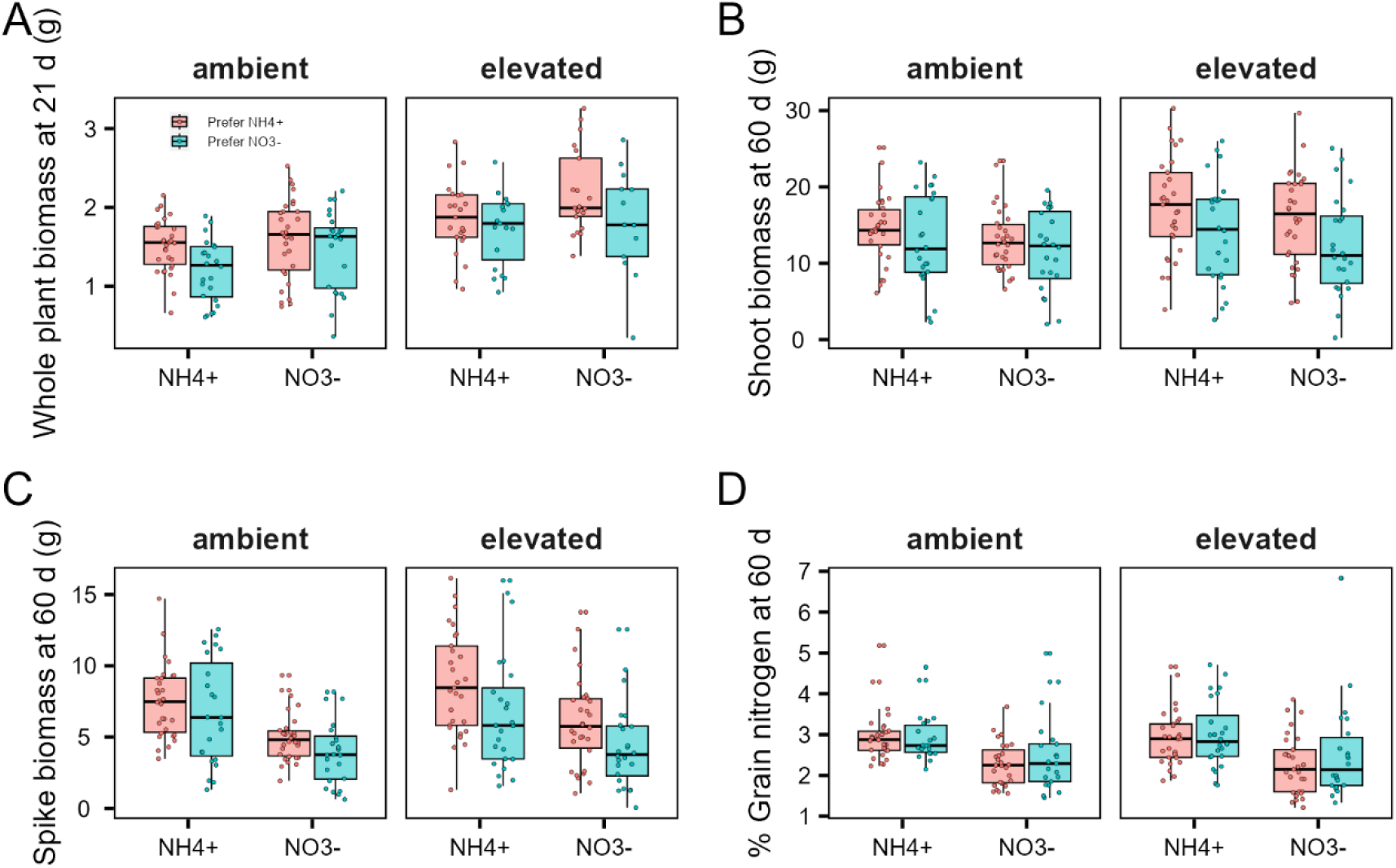
Adaptation to nitrogen forms explained biomass production under both nitrogen forms and CO_2_ levels. The box colors denote classification of nitrogen-form preference at ambient CO_2_ with NH_4_^+^ (pink box) or NO_3_^−^ (blue box)-preferred. Distribution of whole plant biomass at 21 d (a), shoot biomass at 60 d (b), spike biomass at 60 d (c) and developing grain nitrogen concentration at 60 d (d) after transplanting of 9 wheat genotypes under NH_4_^+^ or NO_3_^−^ as a sole nitrogen source in ambient or elevated CO_2_ concentrations. The box plots represent minima, 25^th^ percentiles, medians, 75^th^ percentiles and maxima. Each dot represents each plant replicate across genotypes with the same nitrogen preference classifications.

Relationships among growth traits indicated that several changes across yield components were unique to an individual nitrogen form (Supplementary Figure S11-15). For example, the relationship between tiller fertility and shoot biomass was positive under NH_4_^+^ nutrition, but negative under NO_3_^−^ nutrition (Supplementary Figure S12-13). Furthermore, changes in tiller and rachis number correlated with spike biomass only under NH_4_^+^ nutrition (Supplementary Figure S12). Combined changes in yield components, particularly spike number, proportion of fertile tiller, spikelet number, and spike fertility contributed to the observed variations in spike biomass. Lastly, although biomass traits of genotypes with different NH_4_^+^ tolerance levels did not differ (Supplementary Figure S10), we observed weak but significant correlations, between tolerance levels and some yield components (Supplementary Figure S11).

## Discussion

Shoot carbon assimilation converges with root nitrogen assimilation during biomass production. Changes in atmospheric CO_2_ concentrations likely favor C_3_ species in NH_4_^+^-dominated environments but C_4_ species in NO_3_^−^-dominated environments (Bloom et al., 2012). CO_2_ enrichment lowers plant capacity to use NO_3_^−^ by 16.2% but has a negligible influence on NH_4_^+^ usage (Cheng et al., 2012). Adaptations to such changes includes shifts in species distribution (Bloom et al., 2012) and trade-offs between carbohydrates and proteins to maintain grain quality (Bloom & Plant, 2021). This study highlights the importance of incorporating diverse genetic materials and provides further evidence that increased crop reliance on NH_4_^+^ may mitigate the detrimental effects of rising CO_2_ levels.

Here, genotype accounted for most of the biomass variation (Supplementary Table S1). Because most studies are limited to a few species or genotypes within one species, adaptations to nitrogen forms or tolerance to toxic concentrations have rarely been considered. Genotype-specific responses are likely responsible for contradictory results among studies that used a single genotype to represent a species. For example, the wheat variety Yercora Rojo slowed nitrate assimilation under a free-air CO_2_ enrichment in a potentially high NO_3_^−^ soil (Bloom et al., 2014). In contrast, the variety Batis showed higher nitrogen acquisition in NO_3_^−^-fertilized soil than NH_4_^+^-fertilized soil under elevated daytime CO_2_ levels (Dier et al., 2018). Furthermore, researchers often assume that wheat prefers NO_3_^−^ over NH_4_^+^ or may supply the plants with potentially toxic concentrations of NH_4_^+^. For example, the variety Chuanmai 58 showed significantly slower growth with NH_4_^+^ than NO_3_^−^ at 5 mM (Wang et al., 2020), a concentration 10 times higher than the present study. Our experiment on different genotypes of the same species and near-isogenic lines from the same background allowed us to compare and contrast the genotypic effects. By accounting for potential genetic adaptation to nitrogen forms, we showed that the nitrogen-form preference and NH_4_^+^ tolerance level can explain responses to CO_2_ enrichment (Figure 6).

Preference for a nitrogen form varies with genetic backgrounds and environmental conditions (Britto & Kronzucker, 2013; Miller & Cramer, 2005). Species develop at different rates under NH_4_^+^ or NO_3_^−^ (Bloom et al., 2012). Plants should be adapted to the prevalent nitrogen form in their natural habitats (Gigon & Rorison, 1972; Maathuis, 2009). Soil inorganic nitrogen distribution and relative availability may vary 2 to 3 orders of magnitude (R. B. Jackson & Caldwell, 1993). Microbial activities are responsible for the transient nature of soil nitrogen (L. E. Jackson et al., 1989) that interferes with determining crop nitrogen-form preference based solely on relative availability in the soil. Given the high uptake rates by both plants and microbes, the relative availability may not be a major selective pressure (L. E. Jackson et al., 1989). Perhaps, plants benefit most from being generalists that can employ both forms (Houlton et al., 2007). Despite being statistically significant, nitrogen-form preference may only account for a small proportion of biomass variation (Supplementary Table S2). Plant responses to different nitrogen forms indicate the desirability of including genetically and phenotypically diverse materials when testing for physiological responses that may underlie crop local adaptations to soil nutrients.

Nutrient deficiencies or exposures to high nutrient concentrations usually constrain species distribution range and drive local adaptation (Baxter et al., 2010; Terés et al., 2019). Such adaptations to extreme conditions may diminish overall fitness when these stressors are absent (Mauro & Ghalambor, 2020). Here, after plants were exposed to moderate NH_4_^+^ levels, we observed weak correlations between NH_4_^+^ tolerance and several yield components that suggest potential fitness trade-offs (Supplementary Figure S11). The high light intensity we employed may have mitigated both NH_4_^+^ toxicity (Esteban et al., 2016) and CO_2_ inhibition of NO_3_^−^ photoassimilation (Rubio-Asensio & Bloom, 2016). To evaluate the influence of NH_4_^+^ tolerance on fitness, subsequent investigations should carefully select genetic materials exposed to both high and optimal nitrogen concentrations.

CO_2_ acclimation lowered the biomass stimulation effects of CO_2_ enrichment over time (Bloom et al., 2012). Interestingly, genotypes classified as NH_4_^+^-preferring maintained the biomass stimulation effects after prolonged CO_2_ exposure (Figure 3). Moreover, NH_4_^+^-adapted genotypes also showed higher biomass than NO_3_^−^-adapted genotypes when receiving either form as a sole nitrogen source and at both ambient and elevated CO_2_ levels (Figure 6). These results suggest that plants adapted to NH_4_^+^ may be more resilient to changes in CO_2_ levels (Bloom et al., 2010; Fernando et al., 2017; Subbarao et al., 2021; Subbarao & Searchinger, 2021). Experiments on additional materials should verify whether this finding is universal.

The influence of nitrogen forms on crop development, particularly flowering time, remains unresolved (Kasemsap & Bloom, 2022). Typically, CO_2_ enrichment accelerates development and senescence through rubisco and photosynthesis acclimation (Sicher & Bunce, 1997). Here, nitrogen form had a stronger influence than CO_2_ enrichment as plants transitioned into reproduction, such that biomass responses were similar under both CO_2_ conditions but diverged under different nitrogen forms (Supplementary Table S1 and Figure 4). Regardless of CO_2_ levels, NH_4_^+^ nutrition stimulated flower development and spike biomass accumulation (Figure 4). Several yield components contributed to the increased spike biomass under NH_4_^+^ nutrition (Supplementary Figures 11 & 12). The positive correlations of grain number and spikelet number (Rawson, 1970), and spike biomass (Rivera-Amado et al., 2019; Sierra-Gonzalez et al., 2021; Slafer et al., 2015), allowed us to estimate grain number and potential yield, even though we did not directly measure the grain number and grain yield. Moreover, enhanced flower development under NH_4_^+^ nutrition may have reduced the number of floret abortion in basal spikelets, resulting in relatively higher spike fertility (Supplementary Figure S7). Such action may counteract delayed development, an underlying cause of spikelet abortion (Backhaus et al., 2023). Different nitrogen forms likely triggered a distinct suite of developmental changes (Supplementary Figure S12-13), as previously reported in the model species (Katz et al., 2022).

Improving nitrogen assimilation in elevated CO_2_ atmospheres is critical to human nutrition (Myers et al., 2014) and climate change mitigation (Gao & Cabrera Serrenho, 2023). Wheat may have already sacrificed carbon gain to maintain stable nitrogen concentration as atmospheric CO_2_ levels increased (Bloom & Plant, 2021). Here, adaptations to NH_4_^+^ increased spike mass but did not change grain nitrogen concentrations (Figure 6). Grain nitrogen concentrations in this study are within a range observed in other field settings (Gaju et al., 2014). As such, the ability to maintain protein with no penalty on carbon yield may be advantageous under CO_2_ enrichment. Still, we were unable to assess the nutritional quality of mature grains because of a malfunctioning growth chamber. During advanced grain-filling stages, nitrogen concentrations may continue to change as plants approach senescence (Salgó & Gergely, 2012). Early flowering would most effectively improve yield when coupled with delayed but short senescence to allow longer grain filling time (Xie et al., 2016). Given urea and NH_4_^+^-based fertilizers are already dominant forms in agricultural production (Nishina et al., 2017), information on translocation of assimilated resources to sink organs at maturity would have major implications for food production under climate conditions anticipated in the near future. Breeding for cultivars with improved NH_4_^+^ adaptation and assimilation capacity may not only increase crop climate resilience, but also potentially accelerate development and improve yield without sacrificing nutritional values.

## Supplementary data

Supplementary Table S1. P-values of ANOVA for biomass traits

Supplementary Table S2. P-values of ANOVA for nitrogen preference and NH_4_^+^ tolerance

Supplementary Figure S1 Experimental design.

Supplementary Figure S2 As CO_2_ level increases during the vegetative stage, Veery 10 under NH_4_^+^ nutrition partitioned more biomass to shoot growth.

Supplementary Figure S3 At 60 d of elevated CO_2_ exposure, biomass stimulation slowed down, but CO_2_ still influenced tiller development and seed biomass partitioning under different nitrogen forms.

Supplementary Figure S4 Shoot to root ratio consistent with biomass partitioned to shoot

Supplementary Figure S5 Tiller and spike development

Supplementary Figure S6 Rachis and spikelet development

Supplementary Figure S7 Spike fertility

Supplementary Figure S8 Leaf biomass at 60 d varied with nitrogen forms, CO_2_ levels and genotypes.

Supplementary Figure S9 Nitrogen concentrations of developing grains were generally higher under NH4+ nutrition, except for smaller grains of P10, P33 and P39.

Supplementary Figure S10 The degree of tolerance to high NH_4_^+^ concentration was not associated with biomass production under the environments tested in this study.

Supplementary Figure S11 Correlation of biomass traits

Supplementary Figure S12 Correlation of biomass traits of plants receiving NH_4_^+^

Supplementary Figure S13 Correlation of biomass traits of plants receiving NO_3_^−^

Supplementary Figure S14 Correlation of biomass traits of plants under the ambient CO_2_ concentration

Supplementary Figure S15 Correlation of biomass traits of plants under the elevated CO_2_ concentration

Supplementary Table S3. Growth conditions of previous experiments using wheat cv. Veery 10.

## Acknowledgments

We thanked Joshua Hegarty for discussion on grain measurements. We thanked Teng Vang and the California Wheat Commission for their help on grain protein measurements. We appreciate the assistance of Joshua J. Claxton, Francesco G. Riga, Abenezer D. Shankute, Zennin G. Casl, Steven A. Kunze, Dana R. Lawrence, and James K. Schmidt during data collection.

## Author contributions

Pornpipat Kasemsap and Arnold J. Bloom planned the research and edited the manuscript. Pornpipat Kasemsap collected, analyzed data and wrote the first draft of the manuscript.

## Conflict of Interest

Not applicable.

## Funding

This work was supported in part by USDA-IWYP-16-06702, NSF IOS 13-58675, NSF IOS 16-55810, and the John B. Orr Endowment to Arnold J. Bloom. Pornpipat Kasemsap was supported by Thailand – United States Educational Foundation (TUSEF/Fulbright Thailand), UC Davis Department of Plant Sciences Graduate Student Researcher fellowship, and Henry A. Jastro research award, and Grad Innovator Fellowship from UC Davis Innovation Institute for Food and Health.

## Data availability statement

The data that support the findings of this study are openly available on Dryad at DOI: 10.5061/dryad.00000008p and in supplementary materials. Analysis R codes are available at https://www.github.com/paulkasemsap/Wheat_NxCO2_13gen.

## Notes

### Competing Interest Statement

The authors have declared no competing interest.

## References

Al-Hadeethi, I., Li, Y., Odhafa, A. K. H., Al-Hadeethi, H., Seneweera, S., & Lam, S. K. (2019). Assessment of grain quality in terms of functional group response to elevated [CO _2_], water, and nitrogen using a meta-analysis: Grain protein, zinc, and iron under future climate. Ecology and Evolution, 9(13), 7425–7437. 10.1002/ece3.5210

Asensio, J. S. R., Rachmilevitch, S., & Bloom, A. J. (2015). Responses of Arabidopsis and Wheat to Rising CO _2_ Depend on Nitrogen Source and Nighttime CO _2_ Levels. Plant Physiology, 168(1), 156–163. 10.1104/pp.15.00110

Association of Official Analytical Chemists. (2000). Food composition, additives, natural contaminants (W. Horwitz, Ed.; 17th ed). AOAC International.

Backhaus, A. E., Griffiths, C., Vergara-Cruces, A., Simmonds, J., Lee, R., Morris, R. J., & Uauy, C. (2023). Delayed development of basal spikelets in wheat explains their increased floret abortion and rudimentary nature. Journal of Experimental Botany, erad233. 10.1093/jxb/erad233

Baxter, I., Brazelton, J. N., Yu, D., Huang, Y. S., Lahner, B., Yakubova, E., Li, Y., Bergelson, J., Borevitz, J. O., Nordborg, M., Vitek, O., & Salt, D. E. (2010). A Coastal Cline in Sodium Accumulation in Arabidopsis thaliana Is Driven by Natural Variation of the Sodium Transporter AtHKT1;1. PLoS Genetics, 6(11), e1001193. 10.1371/journal.pgen.1001193

Bishop, D. L., & Bugbee, B. G. (1998). Photosynthetic capacity and dry mass partitioning in dwarf and semi-dwarf wheat (Triticum aestivum L.). Journal of Plant Physiology, 153(5–6), 558–565. 10.1016/S0176-1617(98)80204-6

Blake, N. K., Pumphrey, M., Glover, K., Chao, S., Jordan, K., Jannick, J.-L., Akhunov, E. A., Dubcovsky, J., Bockelman, H., & Talbert, L. E. (2019). Registration of the Triticeae-CAP Spring Wheat Nested Association Mapping Population. Journal of Plant Registrations, 13(2), 294–297. 10.3198/jpr2018.07.0052crmp

Bloom. (2015). The increasing importance of distinguishing among plant nitrogen sources. Current Opinion in Plant Biology, 25, 10–16. 10.1016/j.pbi.2015.03.002

Bloom, A. J., Asensio, J. S. R., Randall, L., Rachmilevitch, S., Cousins, A. B., & Carlisle, E. A. (2012). CO2 enrichment inhibits shoot nitrate assimilation in C3 but not C4 plants and slows growth under nitrate in C3 plants. Ecology, 93(2), 355–367. 10.1890/11-0485.1

Bloom, A. J., Burger, M., Asensio, J. S. R., & Cousins, A. B. (2010). Carbon Dioxide Enrichment Inhibits Nitrate Assimilation in Wheat and Arabidopsis. Science, 328(5980), 899–903. 10.1126/science.1186440

Bloom, A. J., Burger, M., Kimball, B. A., & Pinter, P. J. (2014). Nitrate assimilation is inhibited by elevated CO2 in field-grown wheat. Nature Clim. Change, 4(6), 477–480. 10.1038/nclimate2183 http://www.nature.com/nclimate/journal/v4/n6/abs/nclimate2183.html#supplementary-information

Bloom, A. J., Kasemsap, P., & Rubio-Asensio, J. S. (2020). Rising atmospheric CO _2_ concentration inhibits nitrate assimilation in shoots but enhances it in roots of C _3_ plants. Physiologia Plantarum, 168(4), 963–972. 10.1111/ppl.13040

Bloom, A. J., & Plant, R. E. (2021). Wheat grain yield decreased over the past 35 years, but protein content did not change. Journal of Experimental Botany, 72(20), 6811–6821. 10.1093/jxb/erab343

Bloom, A. J., Smart, D. R., Nguyen, D. T., & Searles, P. S. (2002). Nitrogen assimilation and growth of wheat under elevated carbon dioxide. Proceedings of the National Academy of Sciences, 99(3), 1730–1735. 10.1073/pnas.022627299

Britto, D. T., & Kronzucker, H. J. (2002). NH4+ toxicity in higher plants: A critical review. J Plant Physiol, 159. 10.1078/0176-1617-0774

Britto, D. T., & Kronzucker, H. J. (2013). Ecological significance and complexity of N-source preference in plants. Annals of Botany, 112(6), 957–963. 10.1093/aob/mct157

Britto, D. T., Siddiqi, M. Y., Glass, A. D. M., & Kronzucker, H. J. (2001). Futile transmembrane NH4+ cycling: A cellular hypothesis to explain ammonium toxicity in plants. Proceedings of the National Academy of Sciences, 98(7), 4255–4258. 10.1073/pnas.061034698

Broberg, M., Högy, P., & Pleijel, H. (2017). CO2-Induced Changes in Wheat Grain Composition: Meta-Analysis and Response Functions. Agronomy, 7(2), 32. 10.3390/agronomy7020032

Carlisle, E., Myers, S., Raboy, V., & Bloom, A. (2012). The Effects of Inorganic Nitrogen form and CO(2) Concentration on Wheat Yield and Nutrient Accumulation and Distribution. Frontiers in Plant Science, 3, 195. 10.3389/fpls.2012.00195

Cheng, L., Booker, F. L., Tu, C., Burkey, K. O., Zhou, L., Shew, H. D., Rufty, T. W., & Hu, S. (2012). Arbuscular Mycorrhizal Fungi Increase Organic Carbon Decomposition Under Elevated CO2. Science, 337(6098), 1084–1087. 10.1126/science.1224304

Dier, M., Meinen, R., Erbs, M., Kollhorst, L., Baillie, C.-K., Kaufholdt, D., Kücke, M., Weigel, H.-J., Zörb, C., Hänsch, R., & Manderscheid, R. (2018). Effects of free air carbon dioxide enrichment (FACE) on nitrogen assimilation and growth of winter wheat under nitrate and ammonium fertilization. Global Change Biology, 24(1), e40–e54. 10.1111/gcb.13819

Epstein, E., & Bloom, A. J. (2005). Mineral Nutrition of Plants: Principles and Perspectives. Sinauer Associates, Incorporated.

Esteban, R., Ariz, I., Cruz, C., & Moran, J. F. (2016). Review: Mechanisms of ammonium toxicity and the quest for tolerance. Plant Science, 248, 92–101. 10.1016/j.plantsci.2016.04.008

FAO. (2022, December 13). FAOSTAT: Food Balances (2010-) [Database]. FAOSTAT. https://www.fao.org/faostat/en/#data/FBS

Feng, Z., Rütting, T., Pleijel, H., Wallin, G., Reich, P. B., Kammann, C. I., Newton, P. C. D., Kobayashi, K., Luo, Y., & Uddling, J. (2015). Constraints to nitrogen acquisition of terrestrial plants under elevated CO2. Global Change Biology, 21(8), 3152–3168. 10.1111/gcb.12938

Fernando, N., Hirotsu, N., Panozzo, J., Tausz, M., Norton, R. M., & Seneweera, S. (2017). Lower grain nitrogen content of wheat at elevated CO2 can be improved through post-anthesis NH4+ supplement. Journal of Cereal Science, 74, 79–85. 10.1016/j.jcs.2017.01.009

Fischer, R. A., Moreno Ramos, O. H., Ortiz Monasterio, I., & Sayre, K. D. (2019). Yield response to plant density, row spacing and raised beds in low latitude spring wheat with ample soil resources: An update. Field Crops Research, 232, 95–105. 10.1016/j.fcr.2018.12.011

Fox, J., & Weisberg, S. (2019). An R companion to applied regression (Third edition). SAGE.

Gaju, O., Allard, V., Martre, P., Le Gouis, J., Moreau, D., Bogard, M., Hubbart, S., & Foulkes, M. J. (2014). Nitrogen partitioning and remobilization in relation to leaf senescence, grain yield and grain nitrogen concentration in wheat cultivars. Field Crops Research, 155, 213–223. 10.1016/j.fcr.2013.09.003

Gao, Y., & Cabrera Serrenho, A. (2023). Greenhouse gas emissions from nitrogen fertilizers could be reduced by up to one-fifth of current levels by 2050 with combined interventions. Nature Food, 4(2), 170–178. 10.1038/s43016-023-00698-w

Gifford, R. M., Barrett, D. J., & Lutze, J. L. (2000). The effects of elevated [CO2] on the C:N and C:P mass ratios of plant tissues. Plant and Soil, 224(1), 1–14. 10.1023/A:1004790612630

Gigon, A., & Rorison, I. H. (1972). The Response of Some Ecologically Distinct Plant Species to Nitrate- and to Ammonium-Nitrogen. The Journal of Ecology, 60(1), 93. 10.2307/2258043

Hothorn, T., Bretz, F., & Westfall, P. (2008). Simultaneous Inference in General Parametric Models. 50(3), 346–363.

Houlton, B. Z., Sigman, D. M., Schuur, E. A. G., & Hedin, L. O. (2007). A climate-driven switch in plant nitrogen acquisition within tropical forest communities. Proceedings of the National Academy of Sciences, 104(21), 8902–8906. 10.1073/pnas.0609935104

IPCC. (2021). Climate Change 2021: The Physical Science Basis. Contribution of Working Group I to the Sixth Assessment Report of the Intergovernmental Panel on Climate Change (V. Masson-Delmotte, P. Zhai, A. Pirani, S. L. Connors, C. Péan, S. Berger, N. Caud, Y. Chen, L. Goldfarb, M. I. Gomis, M. Huang, K. Leitzell, E. Lonnoy, J. B. R. Matthews, T. K. Maycock, T. Waterfield, Ö. Yelekçi, R. Yu, & B. Zhou, Eds.). Cambridge University Press. 10.1017/9781009157896

Jackson, L. E., Schimel, J. P., & Firestone, M. K. (1989). Short-term partitioning of ammonium and nitrate between plants and microbes in an annual grassland. Soil Biology and Biochemistry, 21(3), 409–415. 10.1016/0038-0717(89)90152-1

Jackson, R. B., & Caldwell, M. M. (1993). The Scale of Nutrient Heterogeneity Around Individual Plants and Its Quantification with Geostatistics. Ecology, 74(2), 612–614. 10.2307/1939320

Kasemsap, P., & Bloom, A. J. (2022). Breeding for Higher Yields of Wheat and Rice through Modifying Nitrogen Metabolism. Plants, 12(1), 85. 10.3390/plants12010085

Katz, E., Knapp, A., Lensink, M., Keller, C. K., Stefani, J., Li, J.-J., Shane, E., Tuermer-Lee, K., Bloom, A. J., & Kliebenstein, D. J. (2022). Genetic variation underlying differential ammonium and nitrate responses in *Arabidopsis thaliana*. The Plant Cell, 34(12), 4696–4713. 10.1093/plcell/koac279

Langsrud, Ø. (2003). ANOVA for unbalanced data: Use Type II instead of Type III sums of squares. Statistics and Computing, 13(2), 163–167. 10.1023/A:1023260610025

Lenth, R. V. (2023). emmeans: Estimated Marginal Means, aka Least-Squares Means (R package version 1.8.6) [Computer software]. https://CRAN.R-project.org/package=emmeans

Levene, H. (1960). Robust tests for equality of variances. In e a Ingram Olkin (Ed.), Contributions to Probability and Statistic: Essays in honor of Harold Hotelling.

Loladze, I. (2014). Hidden shift of the ionome of plants exposed to elevated CO2 depletes minerals at the base of human nutrition. Elife, 3, e02245.

Maathuis, F. J. (2009). Physiological functions of mineral macronutrients. Current Opinion in Plant Biology, 12(3), 250–258.

Maccaferri, M., Zhang, J., Bulli, P., Abate, Z., Chao, S., Cantu, D., Bossolini, E., Chen, X., Pumphrey, M., & Dubcovsky, J. (2015). A Genome-Wide Association Study of Resistance to Stripe Rust (Puccinia striiformis f. Sp. Tritici) in a Worldwide Collection of Hexaploid Spring Wheat (Triticum aestivum L.). G3: Genes|Genomes|Genetics, 5(3), 449–465. PMC. 10.1534/g3.114.014563

Mauro, A. A., & Ghalambor, C. K. (2020). Trade-offs, Pleiotropy, and Shared Molecular Pathways: A Unified View of Constraints on Adaptation. Integrative and Comparative Biology, 60(2), 332–347. 10.1093/icb/icaa056

Miller, A. J., & Cramer, M. D. (2005). Root nitrogen acquisition and assimilation. In Root Physiology: From Gene to Function (pp. 1–36). Springer, Dordrecht. 10.1007/1-4020-4099-7_1

Myers, S. S., Zanobetti, A., Kloog, I., Huybers, P., Leakey, A. D. B., Bloom, A. J., Carlisle, E., Dietterich, L. H., Fitzgerald, G., Hasegawa, T., Holbrook, N. M., Nelson, R. L., Ottman, M. J., Raboy, V., Sakai, H., Sartor, K. A., Schwartz, J., Seneweera, S., Tausz, M., & Usui, Y. (2014). Increasing CO2 threatens human nutrition. Nature, 510(7503), 139–142. 10.1038/nature13179

Nishina, K., Ito, A., Hanasaki, N., & Hayashi, S. (2017). Reconstruction of spatially detailed global map of NH&lt;sub&gt;4&lt;/sub&gt;&lt;sup&gt;+&lt;/sup&gt; and NO&lt;sub&gt;3&lt;/sub&gt;&lt;sup&gt;−&lt;/sup&gt; application in synthetic nitrogen fertilizer. Earth System Science Data, 9(1), 149–162. 10.5194/essd-9-149-2017

Pleijel, H., Broberg, M. C., Högy, P., & Uddling, J. (2019). Nitrogen application is required to realize wheat yield stimulation by elevated CO _2_ but will not remove the CO _2_ -induced reduction in grain protein concentration. Global Change Biology, 25(5), 1868–1876. 10.1111/gcb.14586

Raun, W. R., & Johnson, G. V. (1999). Improving Nitrogen Use Efficiency for Cereal Production. Agronomy Journal, 91(3), 357–363. 10.2134/agronj1999.00021962009100030001x

Rawson, H. (1970). Spikelet Number, Its Control and Relation to Yield Per Ear in Wheat. Australian Journal of Biological Sciences, 23(1), 1. 10.1071/BI9700001

Rivera-Amado, C., Trujillo-Negrellos, E., Molero, G., Reynolds, M. P., Sylvester-Bradley, R., & Foulkes, M. J. (2019). Optimizing dry-matter partitioning for increased spike growth, grain number and harvest index in spring wheat. Field Crops Research, 240, 154–167. 10.1016/j.fcr.2019.04.016

Royston, J. P. (1982). An Extension of Shapiro and Wilk’s W Test for Normality to Large Samples. Applied Statistics, 31(2), 115. 10.2307/2347973

Rubio-Asensio, J. S., & Bloom, A. J. (2016). Inorganic nitrogen form: A major player in wheat and Arabidopsis responses to elevated CO2. Journal of Experimental Botany, erw465.

Salgó, A., & Gergely, S. (2012). Analysis of wheat grain development using NIR spectroscopy. Journal of Cereal Science, 56(1), 31–38. 10.1016/j.jcs.2012.04.011

Seibert, R., Donath, T. W., Moser, G., Laser, H., Grünhage, L., Schmid, T., & Müller, C. (2021). Effects of long-term CO2 enrichment on forage quality of extensively managed temperate grassland. Agriculture, Ecosystems & Environment, 312, 107347. 10.1016/j.agee.2021.107347

Sicher, R. C., & Bunce, J. A. (1997). Relationship of photosynthetic acclimation to changes of Rubisco activity in field-grown winter wheat and barley during growth in elevated carbon dioxide. Photosynthesis Research, 52(1), 27–38. 10.1023/A:1005874932233

Sierra-Gonzalez, A., Molero, G., Rivera-Amado, C., Babar, M. A., Reynolds, M. P., & Foulkes, M. J. (2021). Exploring genetic diversity for grain partitioning traits to enhance yield in a high biomass spring wheat panel. Field Crops Research, 260, 107979. 10.1016/j.fcr.2020.107979

Slafer, G. A., Elia, M., Savin, R., García, G. A., Terrile, I. I., Ferrante, A., Miralles, D. J., & González, F. G. (2015). Fruiting efficiency: An alternative trait to further rise wheat yield. Food and Energy Security, 4(2), 92–109. 10.1002/fes3.59

Smart, D. R., Ritchie, K., Bloom, A. J., & Bugbee, B. B. (1998). Nitrogen balance for wheat canopies ( *Triticum aestivum* cv. Veery 10) grown under elevated and ambient CO _2_ concentrations. Plant, Cell & Environment, 21(8), 753–763. 10.1046/j.1365-3040.1998.00315.x

Subbarao, G. V., Kishii, M., Bozal-Leorri, A., Ortiz-Monasterio, I., Gao, X., Ibba, M. I., Karwat, H., Gonzalez-Moro, M. B., Gonzalez-Murua, C., Yoshihashi, T., Tobita, S., Kommerell, V., Braun, H.-J., & Iwanaga, M. (2021). Enlisting wild grass genes to combat nitrification in wheat farming: A nature-based solution. Proceedings of the National Academy of Sciences, 118(35), e2106595118. 10.1073/pnas.2106595118

Subbarao, G. V., & Searchinger, T. D. (2021). Opinion: A “more ammonium solution” to mitigate nitrogen pollution and boost crop yields. Proceedings of the National Academy of Sciences, 118(22), e2107576118. 10.1073/pnas.2107576118

Taub, D. R., Miller, B., & Allen, H. (2008). Effects of elevated CO _2_ on the protein concentration of food crops: A meta-analysis: ELEVATED CO _2_ AND CROP PROTEIN CONCENTRATIONS. Global Change Biology, 14(3), 565–575. 10.1111/j.1365-2486.2007.01511.x

Terés, J., Busoms, S., Perez Martín, L., Luís-Villarroya, A., Flis, P., Álvarez-Fernández, A., Tolrà, R., Salt, D. E., & Poschenrieder, C. (2019). Soil carbonate drives local adaptation in *ARABIDOPSIS THALIANA*. Plant, Cell & Environment, 42(8), 2384–2398. 10.1111/pce.13567

Uddling, J., Broberg, M. C., Feng, Z., & Pleijel, H. (2018). Crop quality under rising atmospheric CO 2. Current Opinion in Plant Biology.

Waines, J. G., & Ehdaie, B. (2007). Domestication and Crop Physiology: Roots of Green-Revolution Wheat. Annals of Botany, 100(5), 991–998. 10.1093/aob/mcm180

Wang, F., Gao, J., Yong, J. W. H., Wang, Q., Ma, J., & He, X. (2020). Higher Atmospheric CO2 Levels Favor C3 Plants Over C4 Plants in Utilizing Ammonium as a Nitrogen Source. Frontiers in Plant Science, 11, 537443. 10.3389/fpls.2020.537443

Wujeska-Klause, A., Crous, K. Y., Ghannoum, O., & Ellsworth, D. S. (2019). Lower photorespiration in elevated CO _2_ reduces leaf N concentrations in mature *Eucalyptus* trees in the field. Global Change Biology, 25(4), 1282–1295. 10.1111/gcb.14555

Xie, Q., Mayes, S., & Sparkes, D. L. (2016). Early anthesis and delayed but fast leaf senescence contribute to individual grain dry matter and water accumulation in wheat. Field Crops Research, 187, 24–34. 10.1016/j.fcr.2015.12.009

Xu, G., Fan, X., & Miller, A. J. (2012). Plant Nitrogen Assimilation and Use Efficiency. Annual Review of Plant Biology, 63(1), 153–182. 10.1146/annurev-arplant-042811-105532

Zhang, J., Gizaw, S. A., Bossolini, E., Hegarty, J., Howell, T., Carter, A. H., Akhunov, E., & Dubcovsky, J. (2018). Identification and validation of QTL for grain yield and plant water status under contrasting water treatments in fall-sown spring wheats. Theoretical and Applied Genetics, 131(8), 1741–1759. 10.1007/s00122-018-3111-9

